# Designer DNA NanoGripper

**DOI:** 10.1101/2023.04.26.538490

**Authors:** Lifeng Zhou, Yanyu Xiong, Laura Cooper, Skye Shepherd, Tingjie Song, Abhisek Dwivedy, Lijun Rong, Tong Wang, Brian T. Cunningham, Xing Wang

## Abstract

DNA has shown great biocompatibility, programmable mechanical properties, and structural addressability at the nanometer scale, making it a versatile material for building high precision nanorobotics for biomedical applications. Herein, we present design principle, synthesis, and characterization of a DNA nanorobotic hand, called the “NanoGripper”, that contains a palm and four bendable fingers as inspired by human hands, bird claws, and bacteriophages evolved in nature. Each NanoGripper finger has three phalanges connected by two flexible and rotatable joints that are bendable in response to binding to other entities. Functions of the NanoGripper have been enabled and driven by the interactions between moieties attached to the fingers and their binding partners. We showcase that the NanoGripper can be engineered to interact with and capture various objects with different dimensions, including gold nanoparticles, gold NanoUrchins, and SARS-CoV-2 virions. When carrying multiple DNA aptamer nanoswitches programmed to generate fluorescent signal enhanced on a photonic crystal platform, the NanoGripper functions as a sensitive viral biosensor that detects intact SARS-CoV-2 virions in human saliva with a limit of detection of ∼ 100 copies/mL, providing RT-PCR equivalent sensitivity. Additionally, we use confocal microscopy to visualize how the NanoGripper-aptamer complex can effectively block viral entry into the host cells, indicating the viral inhibition. In summary, we report the design, synthesis, and characterization of a complex nanomachine that can be readily tailored for specific applications. The study highlights a path toward novel, feasible, and efficient solutions for the diagnosis and therapy of other diseases such as HIV and influenza.

**One-sentence summary:** Design, synthesis, characterization, and functional showcase of a human-hand like designer DNA nanobot

## INTRODUCTION

Machine design and manufacture has extended into micro and nanoscopic dimensions with more mobilities and functions through advances in micro- and nanotechnology (*1, 2*). In nature, the most complex nanoscopic machines are comprised of proteins and nucleic acids through bottom-up self-assembly in cellular compartments. These molecular nanomachines utilize nanoscale protein dynamics to achieve defined biological functions such as in protein synthesis by the ribosome (*3*) and cargo transportation along microtubules by kinesin (*4, 5*). Designer nanoscopic robots, called “nanobots”, that can effectively use external energy or stimuli to generate motions and forces to mimic natural complexes have shown great potential for a variety of applications in medicine (*6, 7*). In principle, an effective designer mechanical nanobot for biomedical applications contains rigid compartments that can host and pattern multiple external ligands for target recognition, while also incorporating mobile elements to grant mechanical motions to perform desired functions. At the same time, the structures must be compatible with biological analytes and environments, while being simple to synthesize by self-assembly and be stable during storage and usage.

As the carrier of genetic information for most living organisms, DNA commonly stays in a ∼ 2 nm diameter double helical structure that is comprised of two single strands held together via Watson–Crick base pairing, with 3.4 nm length for each helical turn (*8*). Such physical features make DNA an ideal molecular building block for the construction of nanoscale objects with precisely defined dimensions and curvatures (*9*). DNA nanostructures can serve as biologically compatible and stable molecular pegboards in two-dimensional (2D) or three-dimensional (3D) space to precisely control ligand spacing, valency, and spatial arrangements with nanometer accuracy (*10-22*). Double-stranded DNA (dsDNA) and DNA nanostructures have shown excellent programmable mechanical properties and structural addressability to arrange external ligands (e.g., aptamers, peptides, proteins, nanoparticles) into desired patterns at nanometer scale, making it a versatile material for building nanorobotics employed in various applications (*23, 24*).

In recent years, DNA origami technology(*20*) has proven its ability in the design and fabrication of nanoscale mechanisms and machines such as compliant mechanisms(*25*), paper origami-inspired mechanisms(*26*), and controllable nanomotors(*27*). In many of the prevailing functions, grasp is a powerful ability that a nanobot can apply to perform important functions. However, no previous studies have achieved a nanoscale mechanism made of single origami piece that contains multiple finger-like structures radiating from a central palm to form a gripper-like complex that is capable of collaboratively functioning like human fingers to grab 3D nanoscale entities (e.g., a virus) on one side, as well as to attach the whole complex to a surface on an opposite side for biomedical applications, such as sensing. In this study, we present the design principle, synthesis, and characterization of a designer DNA nanobot, called the DNA “NanoGripper” (or “DNA NG”, “NG” herein), that contains a palm and four-finger-like structures inspired by bird claws, human hands, and bacteriophages evolved in nature. **Scheme-1** illustrates the schematic of NG’s design, means for functionalization, and use in viral sensing and inhibition as a functional showcase. Each of the NG’s fingers contains three phalanges connected by two rotatable joints. The design of the DNA NG robot is highly sophisticated as all the static and moveable components of the NG robot, which contains 12 joints and 13 parts in total, are assembled as a single DNA origami piece with controlled and restricted bending direction through a precise routing of the scaffold and staple strands.

Functions of the NG have been enabled, and motions of the fingers have in turn been driven by the interactions between ligands attached to the fingers’ surface and their binding partners. We show that the NG can be engineered to interact with or capture different nanometer-scale 3D objects with different dimensions, including gold nanoparticles, gold NanoUrchins, and SARS-CoV-2 virions. The NG’s fingers can carry multiple fluorophore-labeled DNA aptamer nanoswitches (*28*) that are programmed to selectively recognize and bind intact SARS-CoV-2 virions. Upon binding with their target, the nanoswitches release a fluorescent signal that is subsequently enhanced when the NG-virus complex is captured on a photonic crystal (PC) surface through DNA hybridization of complementary sequences prepared on the PC and the NG’s palm. Through this approach, the NG successfully functions as a sensitive and specific viral biosensor probe that detects intact SARS-CoV-2 virions in human saliva in < 30 mins with a limit of detection (LOD) of ∼100 genome copies/mL, providing a RT-PCR comparable sensitivity (*29, 30*). Furthermore, we use confocal microscopy imaging to visualize how the same NG aptamer complex can physically block viral entry into the host cells indicating the viral inhibition by the NG. In summary, we report a design principle, synthesis, and characterization of a complex DNA nanobot that can be readily tailored for specific biomedical applications. The NG based platform provides feasible and efficient solutions for the diagnosis and therapy of other diseases such as HIV and influenza viruses. The DNA NG can be also engineered to serve as a drug delivery vehicle like a bacterial contractile injection system reported recently (*31*).

**Scheme 1.**
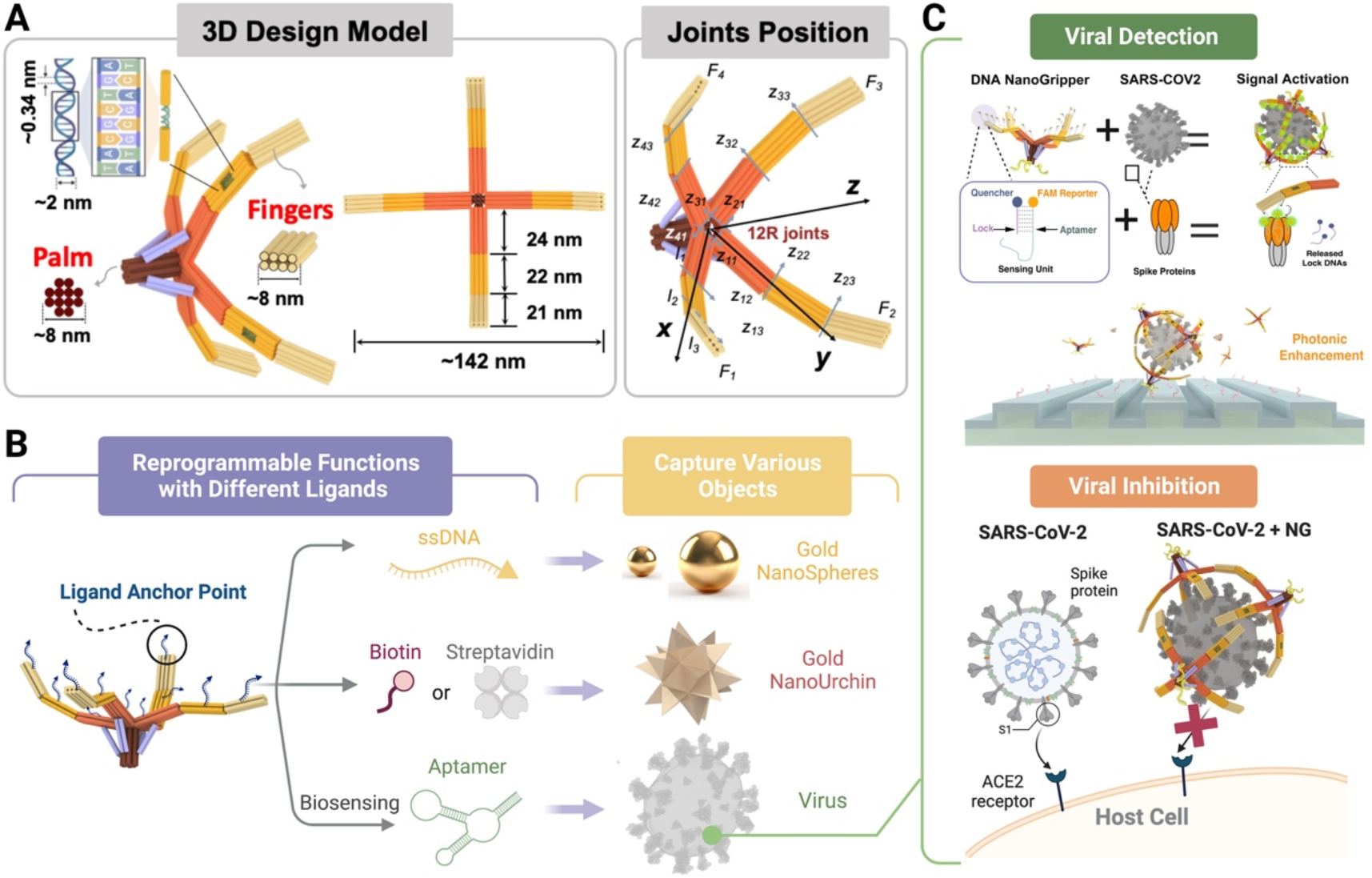
Schematic of the DNA NanoGripper (NG) design, functionalization, and use in viral detection and inhibition. (**A**) Configuration and dimension of the DNA NG that contains a central palm and four bendable fingers, where *F, Z* and *R* represent finger, rotation axis and rotation joint, respectively. (**B**) Functionalization of the DNA NG for interacting with different nanometer scale 3D objects. (**C**) Viral detection an inhibition using DNA NG.

## RESULTS

### DNA NG design principle

Design of the DNA NG was inspired by naturally occurring analogues such as bird claws, human hands, and bacteriophages that have evolved fingers from a central palm domain to effectively grasp objects with various matching geometries and dimensions (**fig. S1A**). Adopting the macroscopic machine design and manufacture procedure, the DNA NG was sculpted, synthesized, and characterized through the steps as illustrated in **fig. S1B**. More specifically in step 1 (nanomachine type synthesis), we decided to make a NanoGripper with four fingers by mainly considering the following two factors: (a) if one finger malfunctions in a three-finger gripper, the remaining two fingers will not function well to grab the targets, and (b) the available scaffold DNA for making DNA origami structures is not long enough to accommodate five-finger design with each finger still containing three phalanges connected by two flexible, human-like fingers, and rotatable joints. In step 2 (nanomachine dimension synthesis), our aim is for the DNA NG to easily grasp objects with overall size of approximate 100 nm diameter or smaller.

The dimensions and parameters of DNA NG design were determined by using the equation and process as detailed in **Note S1**. Accordingly in step 3 (nanomachine design analysis and optimization), we optimized the dimensions of the NG’s palm (12 DNA helices cross section, 60 base pair long) and fingers (8 DNA helices cross section, 200 base pair long) based on the length of the scaffold DNA (8064-base long, see sequence in **Table S1**), as also illustrated in **Fig. 1A** and **fig. S2**. Furthermore, all the fingers are arranged above the end face to make sure that the free DNA NG stays in open configurations (**Fig. 1B**), as fulfilled by incorporating movable hinges on each finger with 4-6 nt long ssDNA connectors based on the rotational orientation of the connected dsDNA helices. In step 4 (determination of the routing of DNA origami scaffold and staples), after the dimensional design of each finger and the connection between four fingers and the palm using 4-6 nt long ssDNA based on the correct rotational orientation of the connected DNA helices, we conducted the entire DNA NG design and the routing of the scaffold DNA using MagicDNA (*32*), oxView (*33*), and caDNAno (*34*), respectively (**Fig. 1C** and **figs. S3-S4**). Additionally, we add dsDNA links between the palm and the fingers to constrain the mobility of the fingers and further ensure that the free DNA NG stays in its open configuration (**fig. S5**). Furthermore, we decorated the inward-facing surface of all four fingers with a total number of 52 ssDNA tethers extending from the staple DNA strands to be used to hybridize with complementary ssDNA or aptamers for enabling the NG’s grabbing motions driven by the interactions with nanometer scale targets. All the staple DNA strands are listed in **Table S2**.

**Fig 1.**
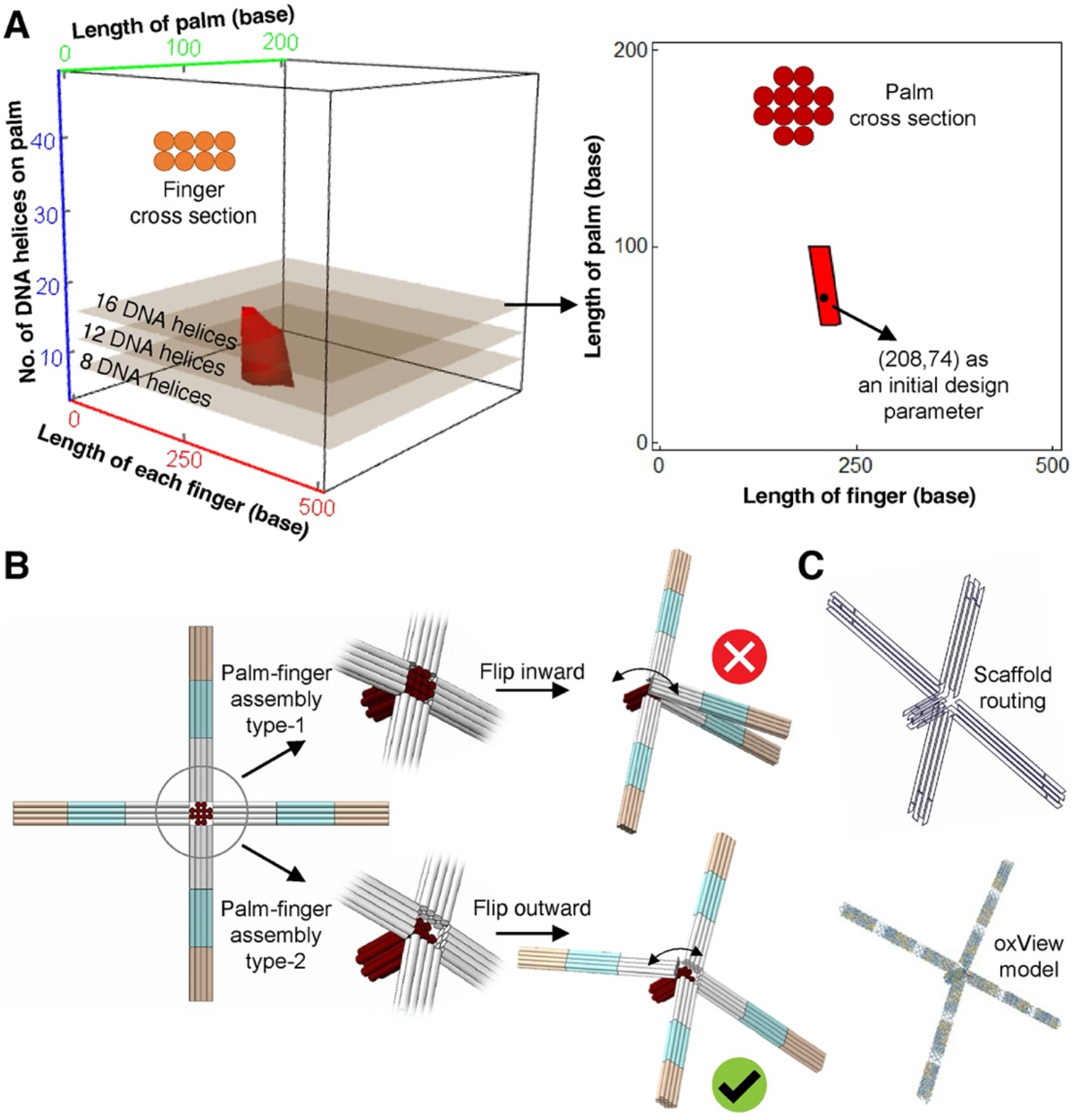
Design of DNA NanoGripper. (**A**) Determination of optimized finger and palm cross-section sizes. (**B**) Schematic of the favorite finger-palm connection design that offers outward flip of the fingers to ensure the NG fingers stay in open configurations in free solution. (**C**) Schematic of scaffold DNA routing displayed using MagicDNA and oxView.

### Synthesis and characterization of DNA NG

The DNA NG is assembled of a long M13mp18 derived “scaffold” DNA (8064-nt long) with 229 short “staple” strands at a 1:10 molar ratio of scaffold:staple after a high-to-low temperature thermal annealing in 1× TAE buffer containing Mg^2+^ ion (**Fig. 2A**). NG formation in the buffers containing different Mg^2+^ concentrations (12 to 22 mM) was optimized and determined as characterized by agarose gel electrophoresis (AGE), which shows that the NG is present as the predominant species in all the Mg^2+^ containing buffers (**Fig. 2B**). However, the NG starts forming higher molecular weight (MW) aggregates that fail to move beyond the wells of the agarose gel, at Mg^2+^ concentration higher than 16 mM. Thus, 1xTAE buffer containing 16 mM Mg^2+^ that provides the needed ionic strength without causing NG aggregation was used to prepare NG assemblies in all subsequent assays. NG structures were visualized by atomic force microscopy (AFM) imaging (**Fig. 2C left** and **fig. S6**) and transmission electron microscopy (TEM) imaging (**Fig. 2C middle** and **fig. S7**) to confirm the successful formation of the NG structure, as projected onto 2D space with different poses that contain four bendable fingers (**Fig. 2D**). Furthermore, cryo-EM images of flash frozen NG samples were collected to show various NG poses in true 3D space (**Fig. 2C right** and **fig. S8**). The NG 3D structure snapshots captured by cryo-EM imaging reveal that, in liquid, all the NGs’ fingers project outward from the central palm to four directions with various phalange joint angles, suggesting the potential of using NGs to grab other 3D objects with matching dimensions.

**Fig. 2.**
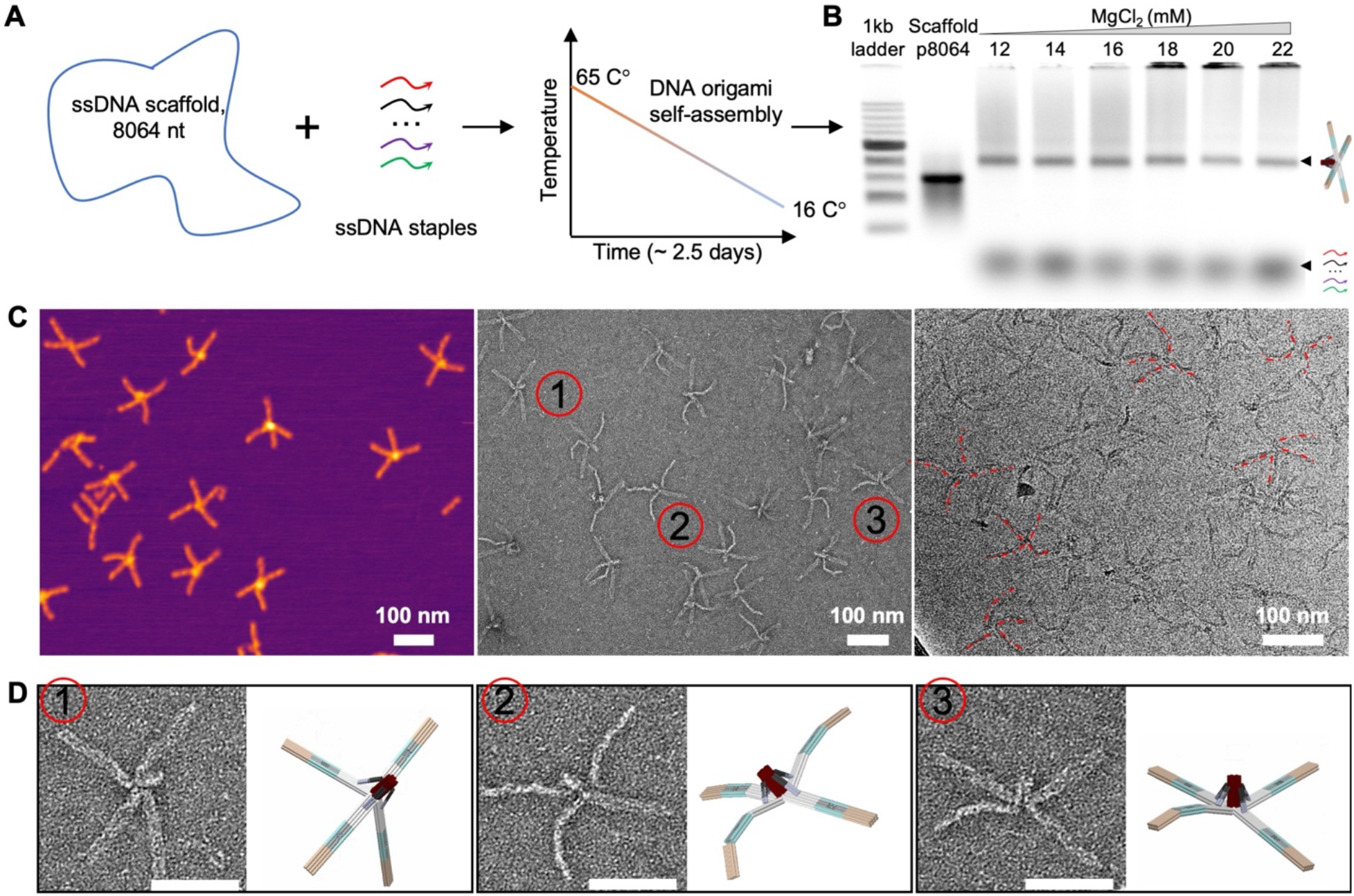
Synthesis and characterization of DNA NanoGripper. (**A**) Schematic of DNA NG assembly using programmed thermal annealing. (**B**) Agarose gel electrophoresis (AGE) shows the NG formation (as a predominate band) in the buffer with different Mg^2+^ concentrations. NG aggregates are observed in higher Mg^2+^ concentration buffers. (**C**) Sample microscopy images of the DNA NG. Left: AFM image, middle: TEM image after negative staining, and right: cryoEM image. (**D**) Side-by-side comparison of DNA NG structure obtained from TEM imaging with the corresponding NG 3D model configurations. Scale bars indicate 100 nm.

### Capture of gold nanoparticle (AuNP) and SARS-CoV-2 virus by DNA NG

The cryo-EM images confirm that the NGs’ fingers can bend like human fingers as expected by our design with various phalange joint angles. We then decorate the NG’s fingers with ssDNA tethers (complementary to the ssDNAs coated on AuNPs) or DNA aptamers (targeting the spike proteins on SARS-CoV-2 outer surface) to demonstrate the potential for using NG to grab 3D objects with matching dimensions. Our AFM and TEM images show that the NG can bend its fingers to accommodate the interaction and capture of spherical-shaped gold nanoparticles (AuNPs) with different sizes (50 or 100 nm diameter), coated with ssDNAs complementary to the ssDNA tethers carried on the NG fingers (**Fig. 3A**). Furthermore, we attached multiple SARS-CoV-2 spike protein-targeting aptamers (*35*) to the NG’s fingers to test its interaction with intact SARS-CoV-2 virions via aptamer-spike interaction. As shown by the cryo-EM images in **Fig. 3B**, multiple NGs can adhere their fingers to the viral particle outer surface with a variety of bending and binding poses due to uneven distributions of spike proteins on SARS-CoV-2 viral outer surface. Some NGs have four fingers grasped on the surfaces while others only have fingertips touching the viral surface. The effective NG-virus interaction further paved the way for utilizing the DNA NG in the scenarios for the detection and inhibition of viral infections as demonstrated hereafter with SARS-CoV-2 virus.

**Fig. 3.**
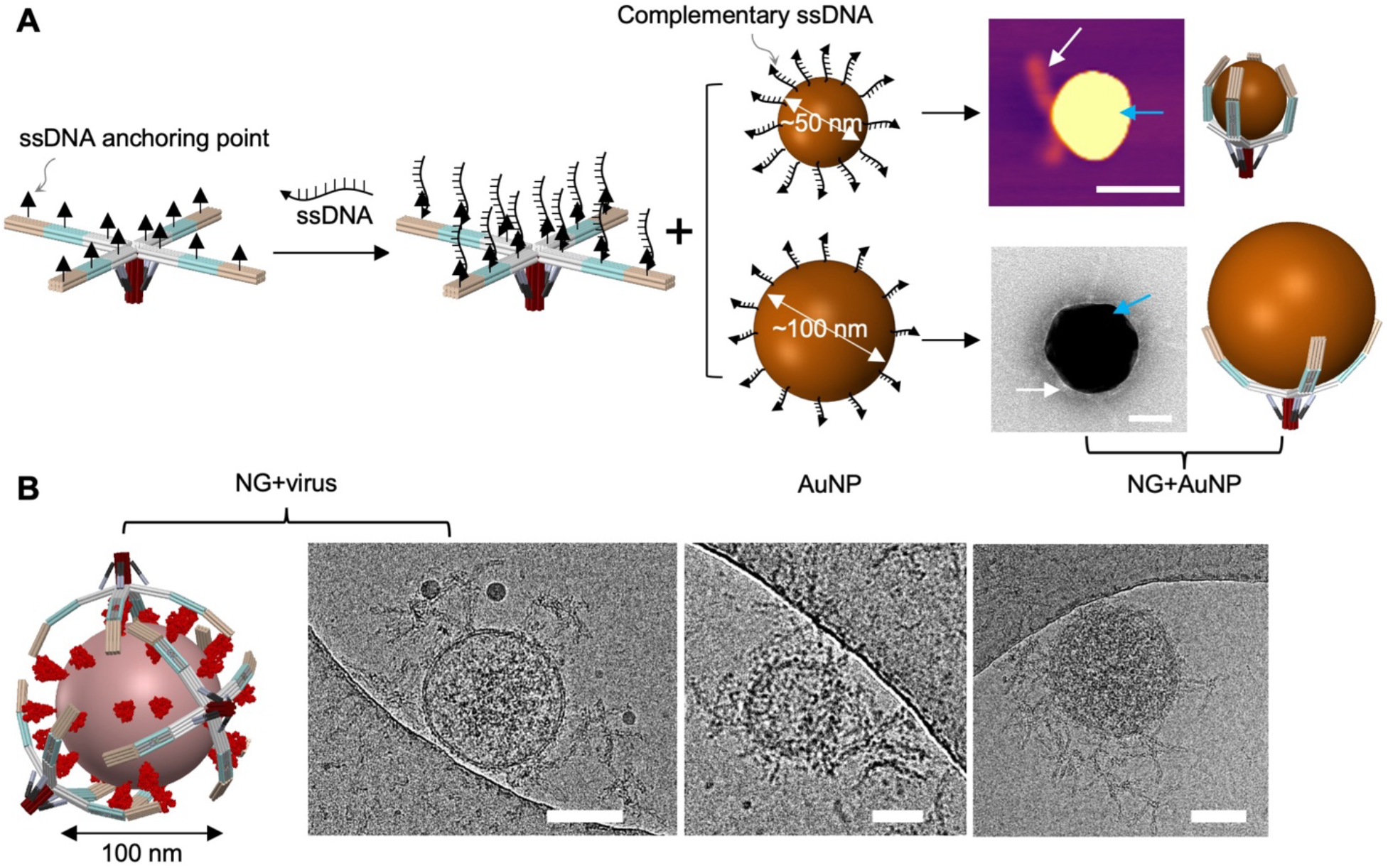
Capture of gold nanoparticles and SARS-CoV-2 virus by DNA NG. (**A**) AFM and TEM images show AuNPs’ interaction with DNA NG carrying ssDNA with a sequence complementary to the ssDNA coated on AuNPs (50 or 100 nm). Scale bars indicate 50 nm. (**B**) Cryo-EM images show the capture of SARS-CoV-2 by the DNA NG carrying multiple spike protein targeting aptamers. Scale bars indicate 50 nm.

### On surface capture of nano-urchin particles (AuNUP) by DNA NG

We have previously demonstrated the ability of the DNA NG to bend its fingers and capture spherical-shaped entities like AuNPs (50 or 100 nm) through DNA hybridization, and SARS-CoV-2 virus (100 nm) via aptamer-protein interaction, in free solution. In this study, we further investigate the interaction between on-surface immobilized DNA NG, functioning as a molecular probe, with its binding partners. To achieve this, we attached five ssDNAs to the bottom of the NG’s palm, which hybridized with the complementary ssDNAs coated on the surface of a photonic crystal (PC) biosensor for surface immobilization and imaging (**Fig. 4A**). Multiple biotinylated ssDNAs were positioned on the NG’s finger surface, facilitating its interaction with streptavidin-coated gold nanourchin particles (AuNUP) to mimic a biological entity carrying surface proteins, captured by the DNA NGs on the PC surface (illustrated in **Fig. 4A**). We employed photonic resonator absorption microscopy (PRAM)(*36*) to visualize surface captured AuNUPs at different concentrations. PRAM has been developed for imaging the attachment of metallic NPs upon a PC surface, enabling the detection of NPs significantly smaller than the diffraction limits. Moreover, metallic AuNPs produce highly localized effects on the PC resonant reflection spectrum, enabling individually attached particles (as small as 30×30×60 nm^3^) to be easily observed using a simple and inexpensive detection instument(*37*). Shown in **Fig. 4B-C**, the DNA NGs immobilized on the PC surface successfully captured AuNUPs from the solution at varying concentrations (0.08, 0.4, 2 pM) as visualized and digitally counted using PRAM at different surface incubation time points (30, 60, 90 mins). This is further illustrated in **Fig. 4D**. Notably, only a very small number of AuNUPs were observed on the bare, non-functionalized PC surface due to nonspecific binding. These results indicate that surface immobilized NGs can effectively capture and detect target entities, such as intact viral particles with high sensitivity, thus expanding the potential applications of the DNA NG for sensing and diagnostics.

**Fig. 4.**
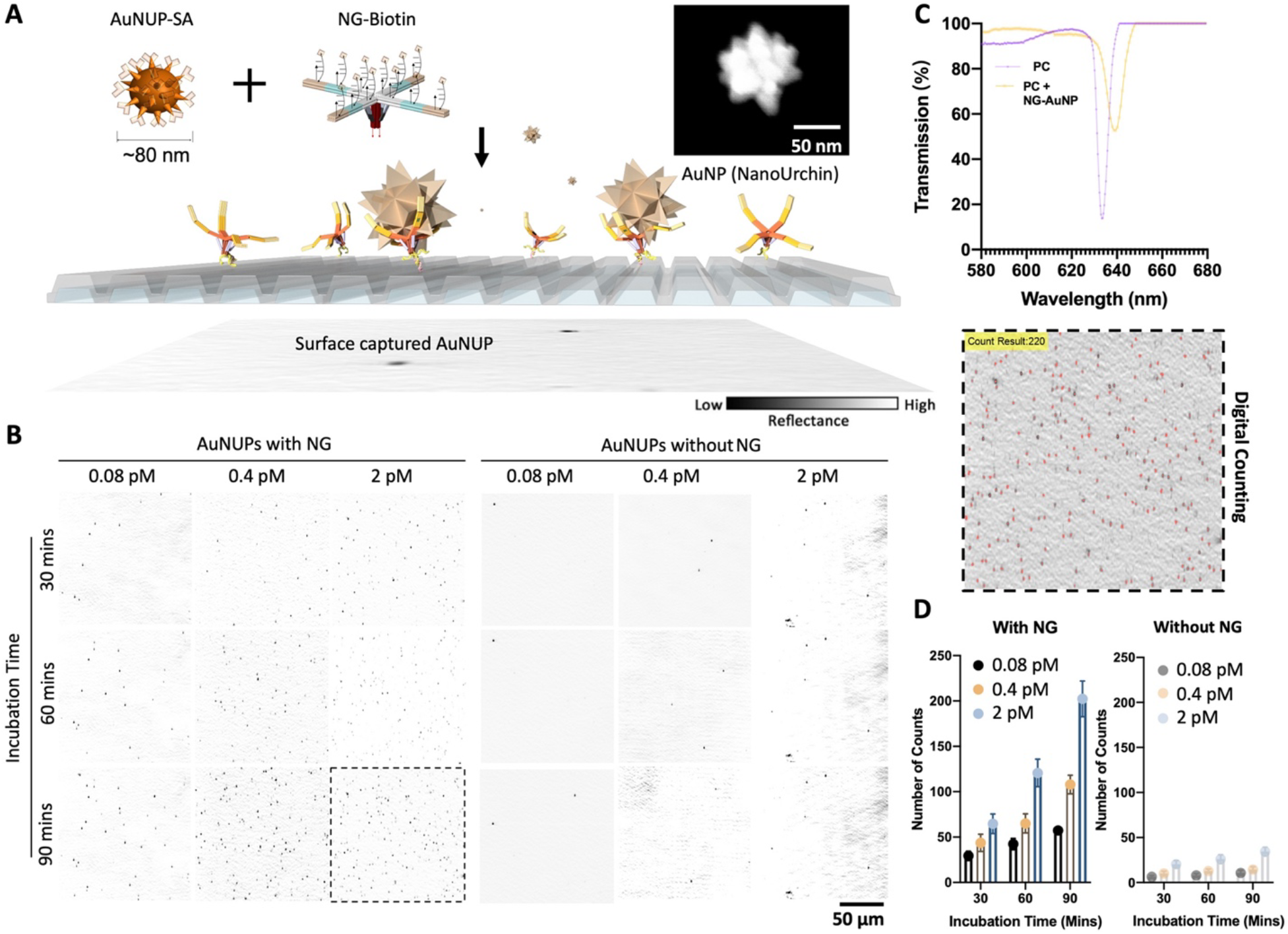
Capture and counting of AuNUPs captured on a PC surface by DNA NGs. (**A**) Schematic of the capture and counting of surface captured AuNUPs by DNA NG. NG is attached to the surface through the hybridization of the ssDNA on the palm (indicated as red arrows) and complementary ssDNA (pink arrows) on the PC surface. NG interacts with AuNUP via biotin-streptavidin (SA) binding. Size and shape of the AuNUP is characterized using SEM imaging. (**B**) PRAM microscopy images of surface captured AuNUPs (black dots) by DNA NGs tested at different particle concentrations and different incubation time points. (**C**) Transmission scan of the PC surface with or without surface immobilized NGs. (**D**) Plots for the counts of surface captured AuNUPs by DNA NGs tested at different concentrations and different incubation time points. Data are presented as mean ± s.d., n = 3 biologically independent samples.

### Detection of SARS-CoV-2 by DNA NG sensor

After determining the ability of surface immobilized DNA NGs to capture and bring the binding targets to a PC surface for quantification via digital counting, we re-engineered the DNA NG to function as a viral biosensor by attaching SARS-CoV-2 spike protein-targeting aptamer nanoswitches on the surface of the NG’s fingers. As illustrated in **Fig. 5A**, each nanoswitch is programmed to comprise a FAM-labeled aptamer that partially hybridizes with a black hole quencher labeled “lock” DNA oligonucleotide to effectively quench the fluorescence in the absence of SARS-CoV-2 virus. The functional NG with aptamer-lock pairs (52 pairs/NG) generates a fluorescent signal only upon capturing a target SARS-CoV-2 virion, because the aptamers anchored on the fingers prefer to bind the spikes on the viral outer surface. Such aptamer-spike binding has been further strengthened through multivalent interactions facilitated by the DNA NG platform, resulting in the separation of the quencher DNA oligo from the aptamer, thus restoring the fluorescence as a detection signal (*28*). The emitted photons from released fluorescence has been subsequently enhanced using a photonic crystal enhanced fluorescence (PCEF) technology (*38*), which provides high signal-to-noise ratio for digitally counting single emitters for ultrasensitive SARS-CoV-2 detection, Importantly, this approach requires only a simple and inexpensive instrument, making it a practical solution for widespread use in biosensing (**Fig. 5B**).

**Fig. 5.**
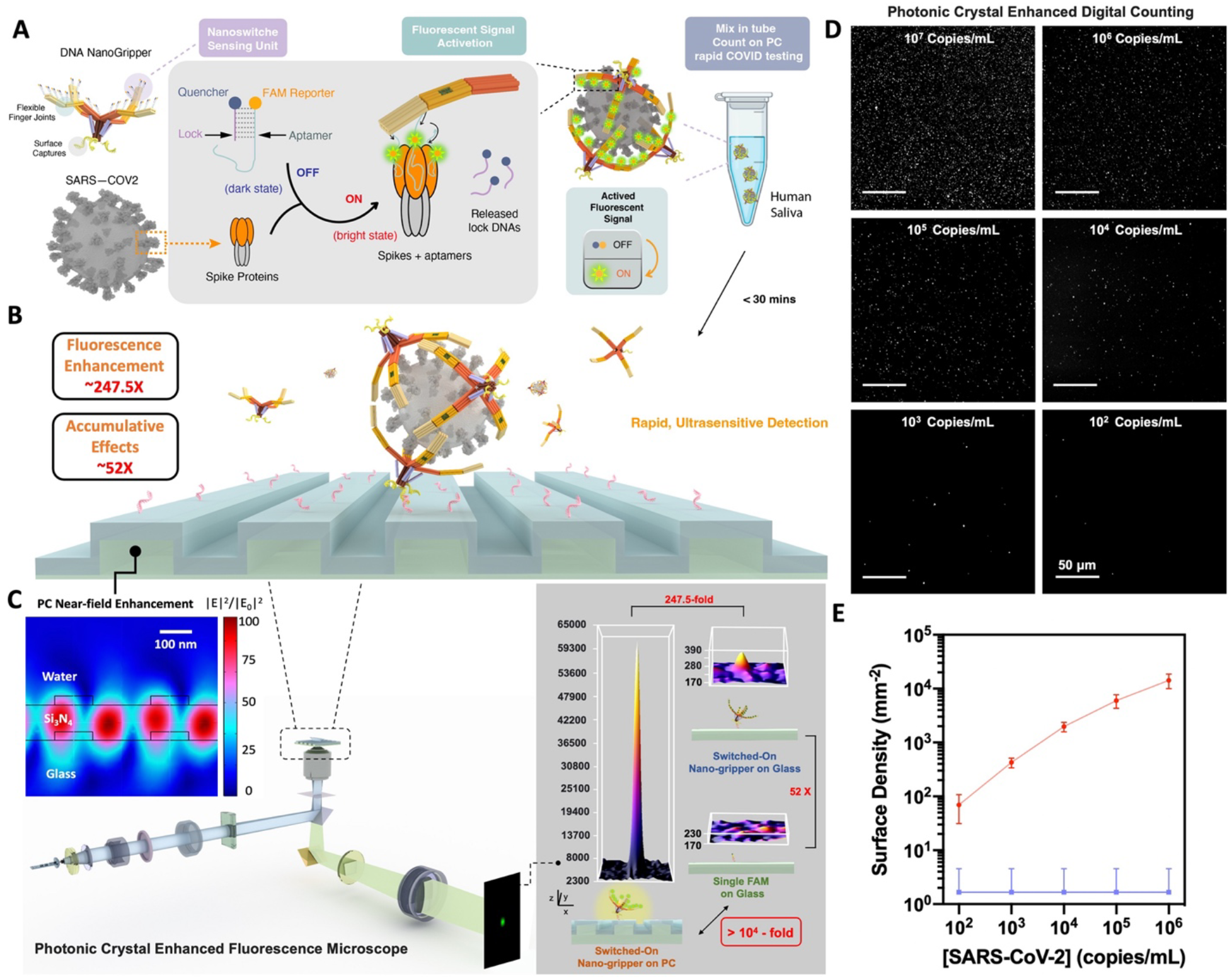
Photonic crystal enhanced detection of SARS-CoV-2 by DNA NG sensor. (**A**) Schematic of viral capture and fluorescent signal generation of functional NGs. (**B**) PC-NG hybrid system. (**C**) PC near-field enhancement and PCEF scanning microscope imaging system with nearly 10^4^-fold signal enhancement compared to an individual FAM reporter on glass substrate. (**D**) PC enhanced digital counting and dose response of SARS-CoV-2 virus detection. Representative scanning images of PC surface captured SARS-CoV-2 at various viral concentrations after 10-min incubation with the NG sensor. (**E**) Quantification of captured SARS-CoV-2 virions on the PC surface at different viral concentrations (red curve). The assays were performed in triplicate with similar observations. Negative control (blue baseline) contains only buffer solution. Error bars represent the standard deviations of three independent measurements.

As dielectric microcavities, PC can provide strong local field enhancement, far-field directional emission, substantial Purcell enhancement, and high quantum efficiency for surface-attached fluorescent reporters (*38-40*). By employing the PC-NG hybrid system, in which the PC enhances fluorescence from unlocked reporters concentrated within captured NG, we report a nearly 10,000-fold signal enhancement compared to a FAM reporter on a glass substrate (**Fig. 5C**). In contrast to previously reported surface-based photonic assays (*41-43*), the capture of SARS-CoV-2 virus by NG sensors occurs rapidly in solution, such that all virions become covered in NGs within 10 mins of mixing. Following the activation of the fluorescent signal by gripped viruses, the virus-NG complex is then incubated on a functionalized PC surface with ssDNA pull-down tethers (via ssDNAs attached to the NG’s palm hybridization with the complementary ssDNA coated on the PC surface) for signal enhancement and digital counting (**Fig. 5D**). Due to the nearly-zero off-target signal achieved with this strategy, we attain a limit of detection (LoD) of 100 genome copies/mL for sensing the intact virus in human saliva (**Fig. 5E**).

### Blocking SARS-CoV-2 viral cell entry by DNA NG

Previous studies have shown 2D or 3D DNA nanostructures can be functionalized to latch onto the viral surface to interrupt viral cell entry by physically blocking viral-cell interactions (*18, 28, 44-47*). Our cryo-EM imaging assays have shown that DNA NG can wrap its fingers around SARS-CoV-2 virus to serve as a physical barrier between the viral particle and a host cell (**Fig. 3B**). Our confocal imaging assays demonstrate that SARS-CoV-2 virions lose their cell internalization ability after being bound by the DNA NG (**Fig. 6**). Membrane-bound structures, including the cell membrane, were stained with DiD (yellow). At the same time, cell nuclei (blue) were stained with Hoechst. SARS-CoV-2 virions were labelled with 6-FAM fluorophore attached to a DNA aptamer that targets the NTD region (under the crown or RBD) of the spike protein on the viral outer surface. Unbound or DNA NG-bound SARS-CoV-2 virions were introduced to the labeled cells. In the unbound condition (**Fig. 6 top row**), SARS-CoV-2 accumulated in the cells over time. In the DNA NG-bound condition (**Fig. 6 bottom row**), SARS-CoV-2 virus accumulation was drastically reduced, and virions were generally electrostatically repelled from cells. Even though some DNA NG-bound virions reached the cell membrane, they were unable to enter cells over a time course of two hours. These results demonstrate the potential of DNA NG as a therapeutic candidate to inhibit viral infections.

**Fig. 6.**
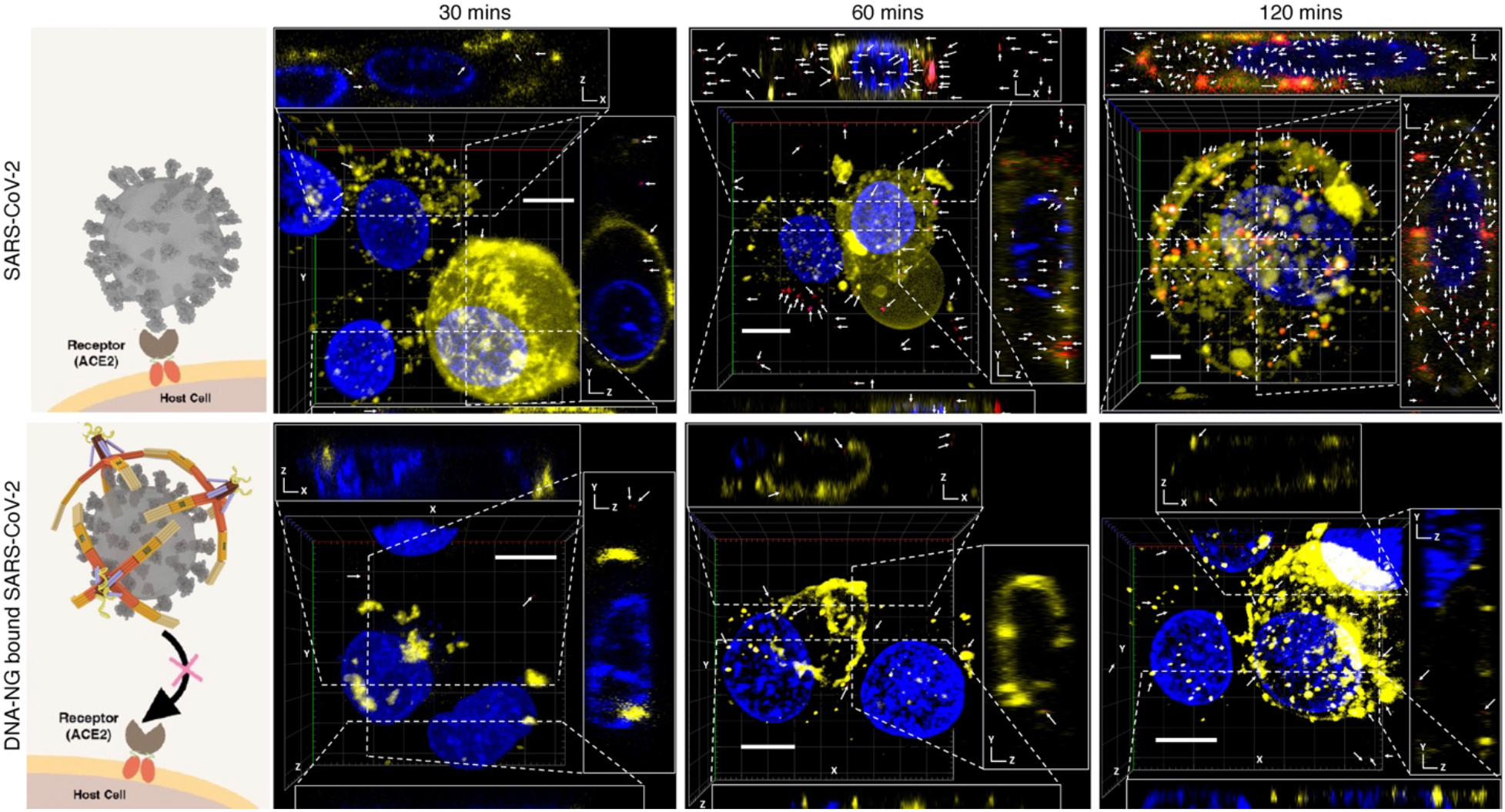
SARS-CoV-2 viral cell entry inhibition by DNA NG. SARS-CoV-2 virions (red), cell membrane structures (yellow), and cell nuclei (blue) were labeled with FAM, Dil and Hoescht stains, respectively. Representative SARS-CoV-2 viral particles are indicated by white arrows. Top row: SARS-CoV-2 virions accumulate over time in the no-inhibitor treated condition. Bottom row: SARS-CoV-2 viral entry is inhibited over time by DNA NG treatment. Each panel is accompanied by three cross-sections of the main view (along the dotted lines in the main images). Confocal images show that DNA NG-unbound SARS-COV2 particles accumulate in the cell (top panels), while DNA NG-bound SARS-COV2 particles are prevented from cell entry (bottom panels). Scale bars indicate 10 µm. Confocal assays were performed in duplicate with similar observations.

## DISCUSSION

Mechanical nanobots have attracted great interest in recent years given their promising potential and suitability for biological and biomedical applications, particularly in disease diagnosis and treatment. In this study, we have successfully adapted the classic mechanical design art and DNA origami technology to develop a design principle and procedure for constructing and functioning a designer DNA NanoGripper (NG) that is inspired by human hands and eagle claws that evolved in nature with a primary function of grabbing. Compared to previous efforts for the design of basic machinery joints/links and simple nanobots using DNA molecules (*23, 24*), the design of the DNA NG robot is much more sophisticated as all the static (palm) and moveable components (fingers), which contains 12 joints and 13 parts in total, are assembled as a single DNA origami piece with controlled and restricted bending direction through a precise routing of the scaffold and staple strands. This new design strategy enables a much more convenient DNA nanobot sample preparation in a one-pot reaction with higher synthetic yield. According to classic machinery category, our NanoGripper can be classified as a parallel open chain mechanism and is one of the most complex DNA nanomachines that have been designed and studied to date. Each NG finger is designed to contain three phalanges connected by two joints, mimicking the function of the human hand to collaboratively promote the grip upon nanometer scale objects with matching dimensions. As a showcase for such capabilities, we attached different target recognition moieties to the NG’s fingers to demonstrate that the NG robot can respectively capture 50 or 100 nm AuNPs (via complementary DNA hybridization), 100 nm AuNUP (via biotin-streptavidin binding), and 100 nm intact SARS-CoV-2 virions (via aptamer-spike protein interaction). Additionally, the NG robot has served as a nanoplatform that carries multiple moieties/ligands to offer higher binding affinity through multivalent interaction, previously demonstrated as an effective means to improve the target binding affinity (*18, 28, 44-47*).

As a possible solution to addressing the challenges for SARS-CoV-2 detection that are typical and historical as evidenced by previous epidemics and pandemics caused especially by emerging RNA viruses for most of the high-impact human viral diseases (*48, 49*), the NG robot fingers are further equipped with multiple nanoswitch motifs to be transformed into a viral sensor that is programmed to release fluorescence signal upon interacting with intact SARS-CoV-2 virions (*28*). This was achieved through the selectivity of SARS-CoV-2 spike protein-targeting aptamer and multivalent spike-aptamer interactions offered by the DNA NG robot platform. In principle, multiple DNA NGs can bind to a single SARS-CoV-2 virus particle resulting in the simultaneous release of many fluorescent reporters from a single virion binding event (**Scheme 1C**). After applying the NG-captured SARS-CoV-2 virion to a photonic crystal surface via hybridizing the ssDNAs on the palm and the ssDNAs anchored on the PC surface, the released fluorescence signal was further enhanced by a microscopy built on photonic crystal enhanced fluorescence (PCEF) technology. The NG viral probe and PCEF combination gave us an LoD of 100 viral genome copies/mL in saliva-containing solution. The range of detection sensitivity of our DNA NG+PCEF biosensor is identical to that obtained by laboratory-based RT-PCR tests, and better than gold standard antigen-based tests. The DNA NG probe has direct access to the intact viruses in the sample. Therefore, our SARS-CoV-2 detection approach circumvents the need to extract RNA materials from the virus particles, RNA purification, reverse transcription, enzymatic amplification, and thermal cycles, resulting in a rapid and sensitive detection assay that can perform isothermally at room temperature. We estimate that the cost of each test is ∼ $1.02 (**Table S3** for the calculation based on the cost of DNA reagents and PC chip) which is much lower than ELISA based antigen tests (∼ $5-10/test) or lateral flow assay (LFA) based antigen tests (∼ $20 per test) (*50*). As nucleic acid tests generate false positive results from the presence of residual viral RNA from degraded viruses in samples (*51, 52*), our approach, which was designed to directly senses SARS-CoV-2 virions, may also address this issue by letting people know when they are no longer infectious and can come out of quarantine.

In addition to detection, the DNA NG-aptamer complex has exhibited therapeutic potential by providing a physical barrier between virus and host cell, as demonstrated by EM imaging as well as *in vitro* SARS-CoV-2 inhibition assay observed using confocal imaging. It is also known that viruses first bind to negatively charged glycosaminoglycans (GAGs) on host cell surface before invasion (*53-57*). Therefore, the DNA NG-aptamer complex not only relies on multivalent interactions for binding SARS-CoV-2 but also electrostatically traps viral particles from the host cell membrane/GAGs through the negative charges of the DNA NG scaffold. The DNA NG platform can be used to deploy aptamers into various arrangements that further increase binding specificity based on viral surface antigen pattern identity. Additionally, the DNA NG can potentially complement the function of neutralizing antibodies (Nabs) by still providing Nabs access to viral surface antigens through the uncovered regions that are otherwise electrostatically shielded from host cells. As demonstrated in previous studies, multivalence is utilized to turn a weak ligand such as oligosaccharides (*53*), or an already strong such as nanobody (*58*) to an stronger binder. Thus, we speculate that the DNA NG robot can provide synergy between multiple ligands and their binding targets to boost the binding affinity even for a strong binder and further improve its viral inhibitory efficacy. In addition to decorating finger’s surface, the figure tips can be functionalized with desired ligands so the DNA NG can be also engineered to serve as a drug delivery nanobot like a bacterial contractile injection system that was recently reported (*31*). It is worth noting that although DNA origami structures have shown biostability *in vivo* (*14, 16*), they can be UV crosslinked (*19*) and/or coated with biocompatible ligands (*13, 59-61*) to further enhance its *in vivo* biostability by increasing resistance to nuclease degradation and/or low salt denaturation (*62-64*).

In summary, by adopting the classic mechanical design art we have developed a design methodology/strategy for creating a highly complex but versatile DNA nanobot in a single DNA origami piece, which can mimic the human hand to grip nanometer scale objects with matching dimensions. Furthermore, we developed a sensitive, single step “direct” viral recognition platform based on the DNA NG structure for the detection of SARS-CoV-2 infections. The same nanobot structure has also shown the potential for viral inhibition. Following the SARS-CoV-2 detection strategy, our DNA NG nanobot can be reengineered to carry respective viral antigen-binding ligands to diagnose and inhibit other viruses that also possess envelope glycoproteins and pose a severe risk to the human health. Moreover, our DNA nanobot design principle can be applied to the sculpting and construction of other DNA nanomachines with desired motions and functions. Such expansion would promote the development and dissemination of DNA nanotechnology in different disciplines, triggering novel and valuable interdisciplinary research topics and directions.

## Supplementary Information

Materials list, Note S1, Figs S1 to S8, Tables S1 to S3, Reference (*65*)

## Acknowledgements

The authors acknowledge the Imaging Facility of Advanced Science Research Center at GC of CUNY, the Electron Microscopy cores at the Materials Research Laboratory (MRL) at UIUC, and the Imaging Cores at the Carl R. Woese Institute for Genomic Biology (IGB) for the technical assistance.

## Author contributions

Conceptualization: LZ, XW, Methodology: LZ, YX, LC, SS, TS, AD, LR, TW, BTC, XW, Supervision: XW, Writing – original draft: LZ, YX, XW, Writing – review & editing: LZ, YX, LC, SS, TS, AD, LR, TW, BTC, XW

